# RNA secondary structures modulate hnRNPH1-mediated alternative splicing of cold-shock protein RBM3

**DOI:** 10.64898/2025.12.18.694734

**Authors:** Shuhan Shen, Lauren McFarlane, Marc-David Ruepp, Julie Qiaojin Lin, Deepak Khuperkar

## Abstract

Alternative splicing is regulated not only by sequence-specific RNA-protein interactions, but also by dynamic RNA secondary structures that remodel under varying cellular conditions. hnRNPH1, a G-rich motif-binding protein, promotes exon skipping across numerous transcripts, yet how RNA structure shapes its regulatory activity remains unclear. Building on our previous work showing that cooler temperature enhances hnRNPH1 binding to RBM3 pre-mRNA and drives poison exon (PE) skipping, we now identify RNA G-quadruplex (rG4) remodelling as the principal *cis*-acting mechanism underlying this temperature-sensitive regulation. Using small-molecule stabilizers and destabilizers, we show that rG4 folding reduces PE skipping, whereas rG4 destabilization enhances hnRNPH1-mediated PE exclusion. These effects are conserved across neuronal and non-neuronal cells. Integrating proteomics, phosphoproteomics, kinase perturbation assays, and candidates emerging from our prior genome-wide CRISPR screen, we further uncover contributions from temperature-sensitive CDC2-like kinase signalling and the RNA helicase DDX5, a known rG4 resolver and an hnRNPH1/hnRNPF co-factor. Together, our findings establish rG4 remodelling as the dominant driver of hnRNPH1-dependent RBM3 splicing, while implicating helicase activity and kinase signalling as modulatory layers that fine-tune this response. More broadly, this work illustrates how environmentally driven RNA structural transitions interface with enzymatic regulators to achieve temperature-responsive control of alternative splicing.

## Introduction

Alternative splicing (AS) is a fundamental post-transcriptional mechanism that enables a single gene to generate multiple transcript isoforms, vastly expanding the proteomic diversity of higher eukaryotes (Blencowe, 2006). This highly regulated process allows cells to dynamically respond to developmental cues, environmental stimuli such as hypothermia, and cellular stressors like hypoxia (Nilsen & Graveley, 2010; Natua et al., 2021; Neumann et al., 2020). The *trans*-acting elements of AS, primarily composed of RNA-binding proteins (RBPs) including the serine/arginine-rich (SR) proteins and heterogeneous nuclear ribonucleoproteins (hnRNPs), recognize specific RNA sequence motifs and structural elements to modulate splice site selection (Fu & Ares, 2014). Among these, hnRNPH1 binds G-rich motifs with sequence specificity and plays a crucial role in regulating exon inclusion and skipping (Caputi & Zahler, 2001; Uren et al., 2016). A key regulatory mechanism involves the inclusion of poison exons (PEs), which are short exons containing premature termination codons (PTCs) that target transcripts for nonsense-mediated mRNA decay (NMD), reducing gene expression. This mechanism has emerged as a conserved strategy for the autoregulation of splicing factors and RBPs and as a molecular switch in response to cellular stress, including temperature fluctuation (Lareau et al., 2007; Ni et al., 2007).

The cold-inducible RNA-binding motif protein 3 (RBM3) is a highly conserved RBP whose expression is robustly induced by hypothermia. As a member of the cold-shock protein family, RBM3 has been implicated in promoting mRNA stability, translation, and synaptic plasticity (Dresios et al., 2005; Peretti et al., 2015). Importantly, RBM3 confers strong neuroprotective effects in models of neurodegeneration, including Alzheimer’s disease and prion disease. In these systems, RBM3 overexpression preserves synaptic function, prevents neuronal loss, and improves behavioural phenotypes, suggesting that it functions as a molecular regulator of hypothermia-induced neuroprotection (Peretti et al., 2015; Zhu et al., 2019).

We and others have shown that RBM3 expression is regulated through temperature-sensitive alternative splicing of a PE, a mechanism conserved across species and cell types (Lin, Khuperkar et al., 2023; Preußner et al., 2023). We previously identified that hnRNPH1 binds to G-rich motifs within RBM3 pre-mRNA to promote PE skipping, thereby enhancing transcript stability and RBM3 protein production at lower temperatures. Notably, hnRNPH1 shows enhanced binding to RBM3 transcripts upon cooling, indicating a temperature-responsive regulatory mechanism (Lin, Khuperkar et al., 2023).

While hnRNPH1 is a primary sequence-specific regulator of RBM3 PE splicing, several temperature-responsive features of the RNA and its regulatory environment suggest additional layers of control. The RBM3 PE contains G-rich elements capable of forming RNA G-quadruplexes (rG4s), whose stability shifts with temperature (Zhang et al., 2025). Given hnRNPH1’s known differential affinity for structured and unstructured RNA elements (Hacht et al., 2014; Haeusler et al., 2014; Neckles et al., 2019), such remodelling may dynamically modulate hnRNPH1–RNA interactions. Moreover, multiple helicases, including members of the DEAD-box family such as DDX5, can remodel rG4 structures and have been shown to cooperate with hnRNP H/F proteins in splicing regulation (Dardenne et al., 2014; Herdy et al., 2018; Wu et al., 2019). Parallel signalling mechanisms, such as phosphorylation by CDC-like kinases (CLKs), have also been implicated in cooling-responsive splice-site selection (Haltenhof et al., 2020), suggesting a broader framework in which structured RNA elements, helicase activity, and kinase signalling collectively integrate environmental temperature cues. Two non-mutually exclusive mechanisms may therefore contribute to temperature-dependent RBM3 PE regulation: 1. RNA structural remodelling, in which cooling alters rG4 folding state and thereby modulates hnRNPH1 accessibility to its binding motifs; and 2. Cooling-induced post-translational modification of hnRNPH1 or other splicing regulators, including potential phosphorylation events that may influence hnRNPH1 function or that of associated cofactors.

Here, we dissect the interplay between RNA structure, phosphorylation, and helicase biology in temperature-dependent regulation of the RBM3 poison exon. By using rG4-stabilizing and -destabilizing ligands, we directly tested the contribution of RNA structural remodelling to hnRNPH1-mediated exon definition. Through CLK inhibition and phospho-mutagenesis of hnRNPH1 RNA recognition motifs, we assessed how kinase signalling influences this process. Finally, motivated by increasing evidence that RNA helicases can remodel structured G-rich elements and cooperate with hnRNP H/F proteins, we extended our analysis to the cooling-responsive helicase DDX5, an established rG4-resolving factor and hnRNPH1 interactor. This integrated approach reveals an expanded regulatory framework linking RNA structure, helicase function, and environmental temperature to RBM3 expression.

## Results

### Stabilization of rG4 structures reduces RBM3 PE skipping

We previously showed that cooling enhances hnRNPH1 binding to RBM3 pre-mRNA and promotes PE skipping, thereby stabilizing the transcript and increasing protein expression (Lin, Khuperkar et al., 2023). To investigate whether *cis*-acting RNA structures contribute to this temperature-sensitive regulation, we focused on RNA G-quadruplexes (rG4s), which are enriched in G-rich motifs within the RBM3 PE, remodel in response to temperature, and are known to influence RBP accessibility. Given hnRNPH1’s preference for unstructured over structured G-rich RNA elements (Martinez-Contreras et al., 2007; Neckles et al., 2019), we reasoned that rG4 remodelling could provide a structural switch for hnRNPH1 recruitment. To test this, iPSC-derived cortical neurons (i3-neurons) were treated with the rG4 stabilizer pyridostatin (PDS), together with SMG1 inhibition to prevent nonsense-mediated mRNA decay (NMD) and enable quantification of PE inclusion. RT-PCR using isoform-specific primers, followed by Percent Spliced In (PSI) analysis, revealed that PDS treatment significantly reduced PE skipping, indicating that stabilization of rG4 structures impairs the exclusion of the PE (Fig. 1A-E). This effect was conserved in HeLa cells, demonstrating that rG4-mediated regulation of RBM3 splicing is not limited to neurons (Fig. 1F-H). These findings are consistent with a recent study in HEK cells that also reported PDS-dependent modulation of RBM3 PE splicing (Zhang et al., 2025), as well as prior work showing that rG4 stabilization reduces hnRNPH1 binding to G-rich RNA (Khateb et al., 2007; Haeusler et al., 2014). Together, these results establish that rG4 folding functions as a *cis*-acting determinant of RBM3 PE regulation, likely by modulating hnRNPH1-RNA interactions.

**Figure 1:**
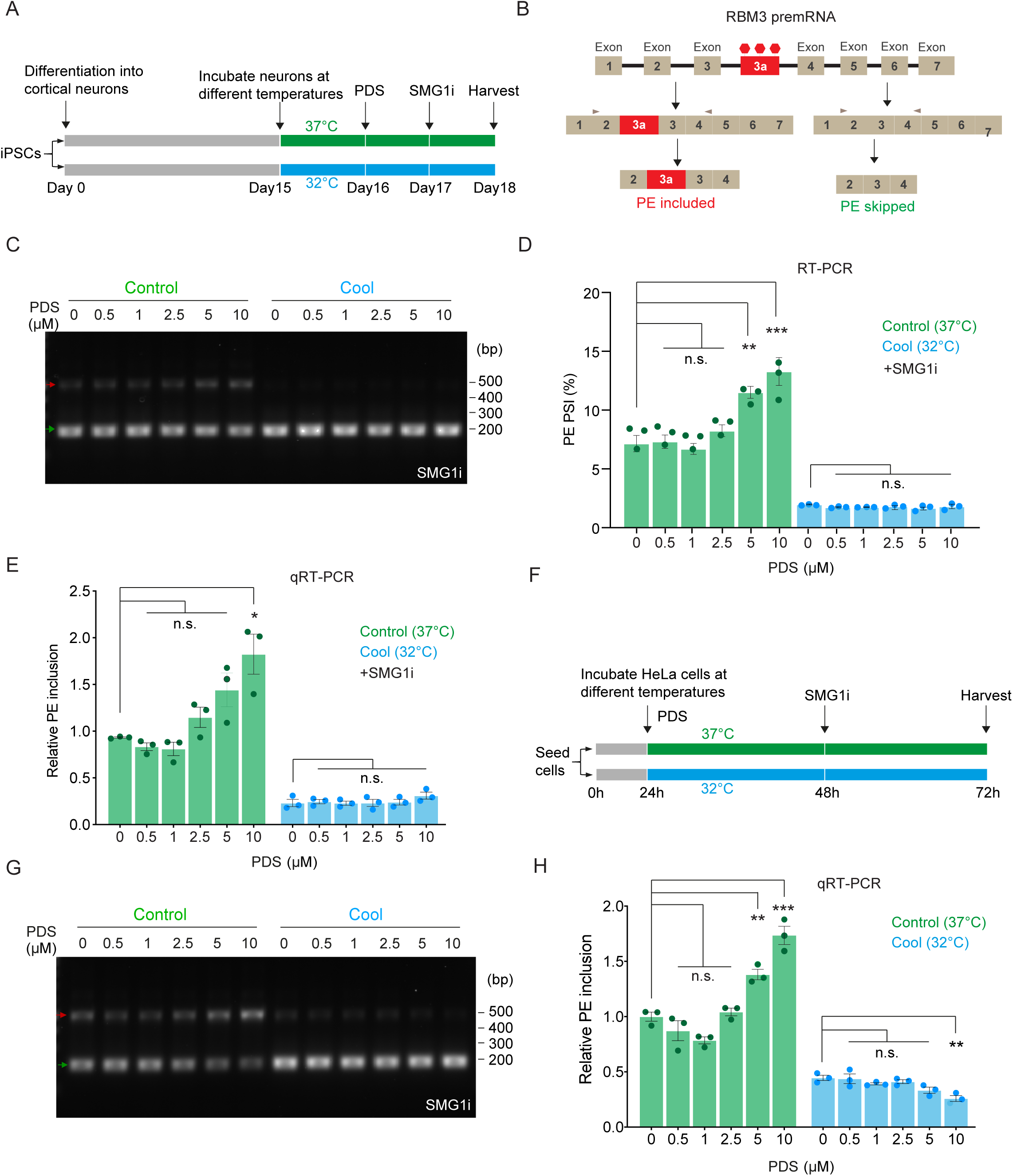
Stabilization of rG4 structures decreases RBM3 PE skipping. See also Figure S1. (A) Schematic of the experimental design in wildtype i3-neurons to evaluate the effect of PDS in RBM3 PE inclusion (B) Schematic of RBM3 Exon 3a, or poison exon (PE), alternative splicing and the resulting PE-included (left) or PE-skipped (right) mRNA products. RT-PCR primer pair amplifying Exon 2–4 are indicated by grey arrows. (C) RT-PCR of RBM3 mRNA (Exon 2–4) in i3-neurons at 37°C or 32°C (72h) in the presence of increasing concentration of PDS (0.5-1.0 µM) showing the PE-included (red arrows) and PE-skipped (green arrow) isoforms. SMG1 inhibitor (SMGi) is used to inhibit NMD. (D) Graph showing the PSI values of RBM3 PE which are calculated based on the intensity of PE-included (red arrows) and PE-skipped (green arrow) isoforms from (C). (E) qRT-PCR quantifying the PSI values of RBM3 PE relative to RBM3 mRNA in i3-neurons at 37°C or 32°C (72h) in the presence of increasing concentration of PDS (0.5-1.0 µM). (F) Schematic of the experimental design in HeLa to evaluate the effect of PDS in RBM3 PE inclusion. (G) RT-PCR of RBM3 mRNA (Exon 2–4) in HeLa at 37°C or 32°C (72h) in the presence of increasing concentration of PDS (0.5-1.0 µM) showing the PE-included (red arrows) and PE-skipped (green arrow) isoforms. (H) qRT-PCR quantifying the PSI values of RBM3 PE relative to RBM3 mRNA in HeLa at 37°C or 32°C (72h) in the presence of increasing concentration of PDS (0.5-1.0 µM). Data information: N = 3 biological replicates. Mean ± SEM; ns (not significant), *(P<0.05), **(P<0.01); ***(P<0.001); one-way ANOVA with multiple comparisons.

### RBM3 PE skipping is enhanced by destabilization of rG4 structures

To further assess the role of rG4 structures in RBM3 splicing regulation, i3-neurons were treated with the rG4-destabilizing agent TMPyP4 (Huang et al., 2017), followed by SMG1 inhibition. RT-PCR analysis demonstrated that TMPyP4 treatment led to a significant increase in PE skipping, compared to control conditions (Fig. 2), consistent with a model in which rG4 destabilization facilitates exon exclusion. These findings, taken together with the effects of rG4 stabilization, support a model in which structural remodelling of rG4 elements dynamically regulates hnRNPH1-mediated RBM3 splicing.

**Figure 2:**
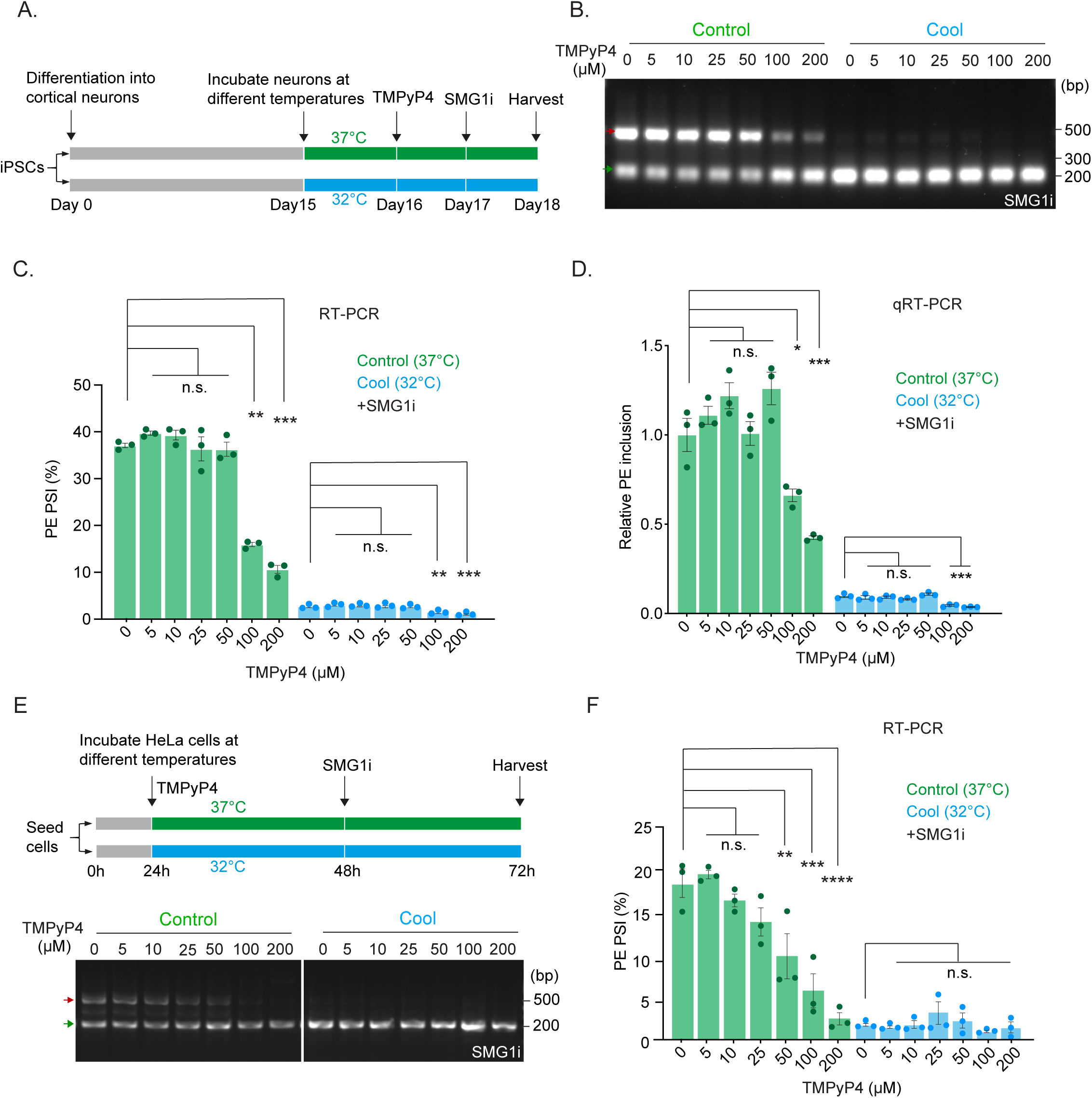
RBM3 PE skipping increases on destabilization of rG4 structures. See also Figure S1. (A) Schematic of the experimental design in i3-neurons to evaluate the effect of TMPyP4 in RBM3 PE inclusion (B) RT-PCR of RBM3 mRNA (Exon 2–4) in i3-neurons at 37°C or 32°C (72h) in the presence of increasing concentration of TMPyP4 (5-200 µM) showing the PE-included (red arrows) and PE-skipped (green arrow) isoforms. (C) Graph showing the PSI values of RBM3 PE which are calculated based on the intensity of PE-included (red arrows) and PE-skipped (green arrow) isoforms from (B). (D) qRT-PCR quantifying the PSI values of RBM3 PE relative to RBM3 mRNA in i3-neurons at 37°C or 32°C (72h) in the presence of increasing concentration of TMPyP4 (5-200 µM). (E) Schematic of the experimental design in HeLa to evaluate the effect of TMPyP4 in RBM3 PE inclusion (top). RT-PCR of RBM3 mRNA (Exon 2–4) in HeLa at 37°C or 32°C (72h) in the presence of increasing concentration of TMPyP4 (5-200 µM) showing the PE-included (red arrows) and PE-skipped (green arrow) isoforms (bottom). (F) Graph showing the PSI values of RBM3 PE which are calculated based on the intensity of PE-included (red arrows) and PE-skipped (green arrow) isoforms from (E). Data information: N = 3 biological replicates. Mean ± SEM; ns (not significant), *(P<0.05), **(P<0.01); ***(P<0.001); one-way ANOVA with multiple comparisons.

### RBM3 PE skipping is regulated by temperature-sensitive CDC2-like kinase (CLK)

We next explored whether post-translational modifications (PTMs) contribute to the differential temperature-dependent hnRNPH1-RBM3 mRNA interaction and hence modulate RBM3 AS. CDC2-like kinases (CLKs) emerged as compelling candidates, as they are known to phosphorylate SR proteins and modulate splicing activity in a temperature-sensitive manner, with increased activity under mild hypothermia (Muraki et al., 2004; Haltenhof et al., 2020). To test the functional relevance of CLK activity in RBM3 splicing, we treated cells with the CLK inhibitor TG003 and assessed RBM3 PE inclusion via RT-PCR. CLK inhibition led to increased PE inclusion (Fig. 3A-C), implying temperature-sensitive phosphorylation events in RBM3 splicing regulation. To evaluate whether hnRNPH1 is directly regulated by phosphorylation, we used PhosphoSitePlus (Hornbeck et al., 2015) to predict high-confidence phospho-sites. Selection was based on (a) coverage of all three RNA recognition motifs (RRMs), (b) conservation between human and mouse, and (c) high phosphorylation prediction scores. A panel of phospho-mutant hnRNPH1 constructs was generated and tested using a previously described RBM3 minigene assay (Fig. S2) (Lin, Khuperkar et al., 2023). Percent Spliced in (PSI) analyses from the RT-PCR data revealed that none of the individual phospho-site mutants significantly altered RBM3 PE skipping activity. These findings suggest that phosphorylation at these individual residues is either not essential for splicing regulation or functionally redundant across sites. Alternatively, other *trans*-acting splicing factors, such as SR proteins, may serve as key phosphorylation-dependent regulators in this system.

**Figure 3.**
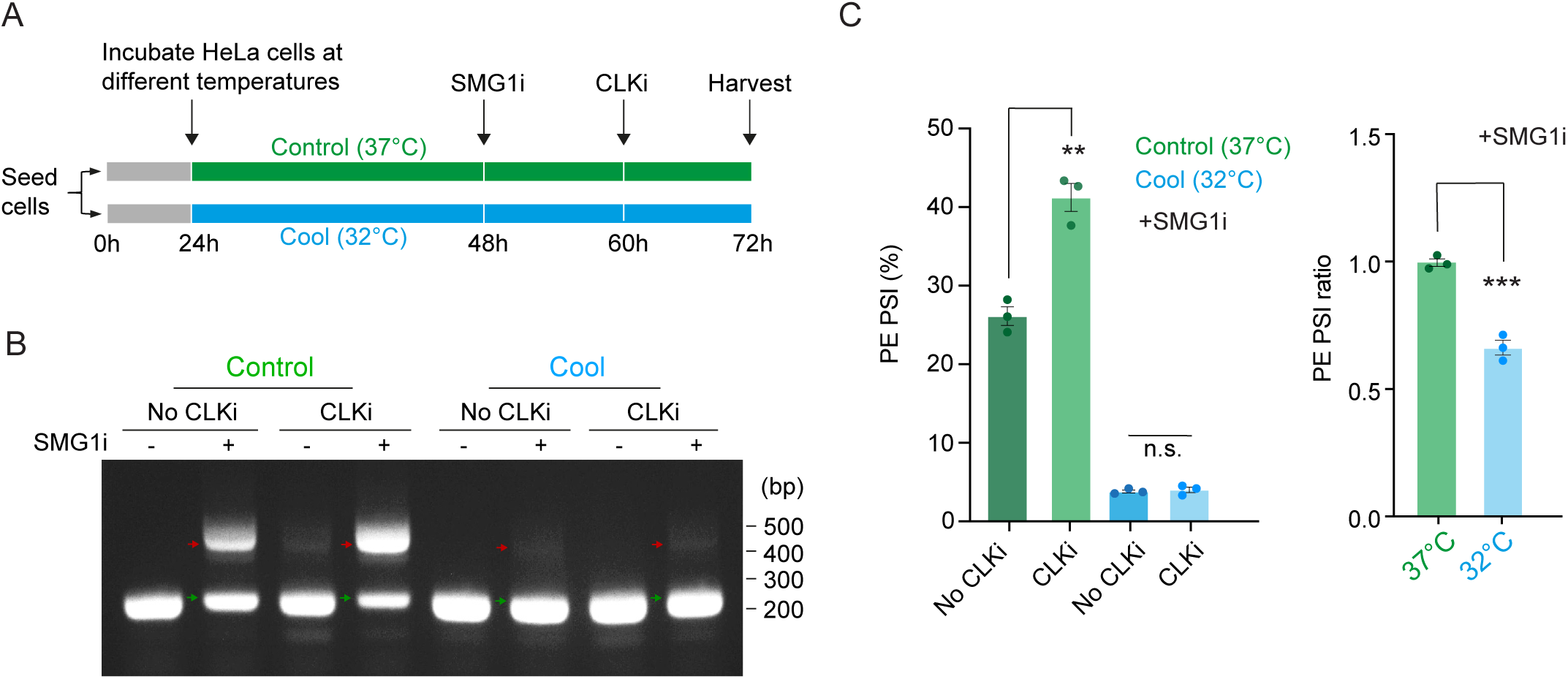
RBM3 PE skipping is regulated by temperature-sensitive kinase CLK. See also Figure S2. (A) Schematic of experimental procedure using CLK inhibitor (CLKi) for evaluating RBM3 PE inclusion. (B) RT-PCR of RBM3 mRNA (Exon 2–4) in HeLa at 37°C or 32°C (72h) in the presence or absence of SMG1 inhibitor and CLK inhibitor showing the PE-included (red arrows) and PE-skipped (green arrow) isoforms. (C) Graph showing the PSI values of RBM3 PE which are calculated based on the intensity of PE-included (red arrows) and PE-skipped (green arrow) isoforms from (B). Data information: N = 3 biological replicates. Mean ± SEM; ns (not significant), *(P<0.05), **(P<0.01); ***(P<0.001); Unpaired t-test in (C).

### DDX5 is a cooling-responsive helicase linking rG4 remodelling to RBM3 PE regulation

From our analyses of RNA structure stability, we next asked whether RNA helicases, which can resolve secondary structures including rG4s, participate in RBM3 PE regulation. To address this, we revisited our previous genome-wide CRISPR knockout screen in i3-neurons (Lin, Khuperkar et al., 2023) and identified three helicases within the top 200 positive regulators of RBM3 expression: DDX58 (P = 0.0070, rank 102), DDX24 (P= 0.0198, rank 215), and DDX5 (P = 0.0203, rank 219) (Appendix Table S3) . Among these, DDX5 emerged as the most compelling candidate because of the following reasons. First, multiple rG4-interactome studies identify DDX5 as one of the strongest and most consistent rG4-binding and rG4-resolving helicases, outperforming other DEAD-box helicases such as DDX3X and DDX21 (Herdy et al., 2018). Second, DDX5 is a known physical interactor of hnRNPH1/hnRNPF and cooperates with hnRNPH1/hnRNPF proteins to regulate alternative splicing structures (Dardenne et al., 2014; Herdy et al., 2018; Wu et al., 2019). Given this convergence, we examined whether DDX5 is temperature responsive. Western blot analyses across multiple cell types—including i3-neurons, HeLa and SH-SY5Y (neuroblastoma cells)—revealed a robust induction (Fold change: 2.19, 1.70 and 1.69-fold respectively) of DDX5 protein levels upon cooling to 32°C for 72 h (Fig. 4A, B, S3A). This was further supported by quantitative proteomics from iPSCs revealing elevated DDX5 abundance (∼1.5 fold), but not DDX17, at 32°C compared to 37°C, showing that DDX5 is a cooling-responsive factor (Fig. 4C). Importantly, phosphoproteomic analysis demonstrated that DDX5 phosphopeptides are significantly enriched at 32 °C, mapping predominantly to RGG/RG-rich regions associated with RNA structure recognition and helicase cofactor interactions, indicating that DDX5 is subjected to cooling-induced post-translational modification (Fig. 4D, S3B). Furthermore, kinase–substrate prediction analysis identified DDX5 as a putative CLK target, with multiple phosphosite-containing peptides ranking highly for CLK family kinases. Notably, two DDX5 peptides scored highest for CLK3 (Fig. S3C) (Johnson et al., 2023). Because CLK inhibition increases RBM3 PE inclusion (Fig. 3), these predictions provide a mechanistic link between cooling-dependent kinase signalling and helicase remodelling. Collectively, these findings position DDX5 as a cooling-regulated rG4-associated helicase, potentially acting downstream of temperature-sensitive kinase signalling to modulate hnRNPH1-dependent RBM3 splicing.

**Figure 4.**
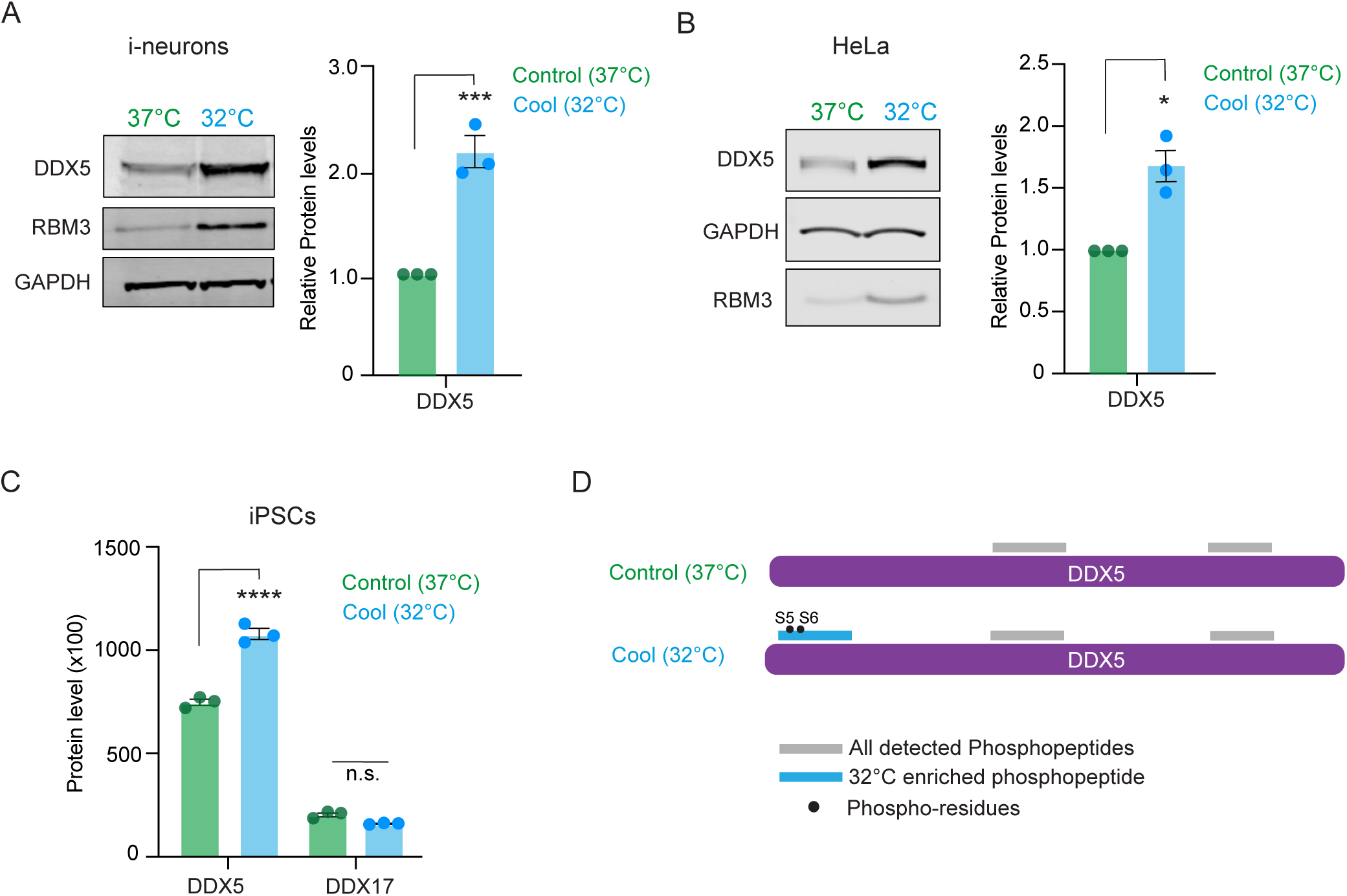
RNA Helicase DDX5 is regulated in a cooling-dependent manner. See also Figure S3. (A) Western blot analysis of DDX5 protein levels in i3-neurons cultured at 37°C or 32°C (72h). Representative blots are shown (left) with quantification normalized to GAPDH (right). (B) Western blot analysis of DDX5 protein levels in HeLa cultured at 37°C or 32°C (72h). Representative blots are shown (left) with quantification normalized to GAPDH (right). (C) Global proteomics analysis from iPSCs grown at 37°C or 32°C showing significantly elevated DDX5, but not DDX17, peptide abundance under cooling conditions. (D) Schematic showing DDX5 phosphopeptides identified from phosphoproteomics analysis. Significantly enriched peptide at 32°C is indicated in blue, with phosphorylated serine residues highlighted. Cooling-responsive phosphosites cluster within RG/RGG-rich regions implicated in RNA structure recognition. Data information: N = 3 biological replicates. Mean ± SEM; ns (not significant), *(P<0.05), **(P<0.01); ***(P<0.001); Unpaired t-test in (A) and Welch’s t-test in (B)

## Discussion

Our study uncovers a central role for RNA G-quadruplex (rG4) structures in regulating temperature-sensitive alternative splicing of the neuroprotective cold-shock protein RBM3. While we showed previously that hnRNPH1 mediates RBM3 poison exon (PE) skipping through binding to G-rich motifs (Lin, Khuperkar et al., 2023), we now demonstrate that the RNA’s structural state significantly influences this regulation. Stabilization of rG4s using pyridostatin impaired PE skipping, whereas rG4 destabilization with TMPyP4 enhanced PE exclusion, identifying rG4s as key *cis*-acting regulatory elements whose folding state influences RBM3 PE splicing.

Our findings build directly on our earlier demonstration that hnRNPH1 binding to RBM3 transcripts is enhanced upon cooling, driving PE skipping and stabilizing RBM3 mRNA (Lin, Khuperkar et al., 2023). These findings refine our earlier model by adding a structural dimension: rG4 unfolding at lower temperatures likely increases accessibility of G-rich motifs to hnRNPH1, thereby promoting PE exclusion. Although we did not directly quantify hnRNPH1-RNA binding under rG4 stabilization/ destabilization, the splicing outcomes align with hnRNPH1’s established preference for unstructured G-tracts. Our results complement recent work showing that many cellular rG4s destabilize upon cooling and act as RNA thermosensors (Zhang et al., 2025). Whereas Zhang et al. highlighted transcriptome-wide consequences of rG4 remodelling, our work provides a mechanistic explanation for how rG4 dynamics translate into altered exon choice, positioning hnRNPH1 as the *trans*-acting player of this structural switch.

In addition to RNA structural changes, our data support a modulatory role for temperature-dependent post-translational modification. Inhibition of CDC2-like kinases (CLKs) increased RBM3 PE inclusion, consistent with prior evidence that CLK activity is enhanced under mild hypothermia (Haltenhof et al., 2020). However, individual phospho-site mutations in hnRNPH1 did not alter splicing outcomes (Fig. 3, S2), suggesting that phosphorylation of hnRNPH1 alone is insufficient to explain temperature-dependent regulation. This raises the possibility of combinatorial phosphorylation, redundancy among sites, or contributions from other CLK-sensitive splicing factors, such as SR proteins.

Notably, the effects of pharmacological rG4 perturbation differed depending on temperature. While rG4 stabilization by pyridostatin strongly reduced poison exon skipping at 37°C, it had no significant effect at 32°C, whereas the rG4-destabilizing ligand TMPyP4 retained a modest effect at 32°C. One possible explanation is that cooling itself induces rapid rG4 unfolding, potentially aided by increased helicase activity, such that exogenous stabilization becomes ineffective once this structural transition has occurred. These observations suggest that rG4 remodelling is an early and dominant event in the cooling response, and that additional regulatory factors may act to reinforce or sustain this unfolded state.

A major conceptual advance of the current study is the identification of DDX5 as a cooling-responsive helicase that intersects with rG4 remodelling and hnRNPH1-mediated splicing. DDX5 emerged as a positive regulator of RBM3 expression in our prior genome-wide CRISPR knockout screen in i3-neurons (Lin, Khuperkar et al., 2023). Here, we show that DDX5 protein levels increase robustly across multiple cell types upon cooling, indicating temperature-responsive regulation. Importantly, DDX5 is a well-established rG4 resolver (Herdy et al., 2018; Wu et al., 2019) and cooperates with hnRNPH1/hnRNPF in G-rich alternative splicing programs (Dardenne et al., 2014). Phosphoproteomics further revealed multiple cooling-induced DDX5 phosphopeptides. While we could not assign these sites directly to CLK activity, DDX5 is a known substrate of SR-related kinases, supporting a model in which cooling modulates helicase function through phosphorylation. Together, these observations place DDX5 at a central intersection between RNA structural remodelling and temperature-dependent signalling, acting as an auxiliary factor that may enhance hnRNPH1 access to rG4-containing regions. Future work should determine the precise biochemical consequences of DDX5 phosphorylation on rG4 unwinding and hnRNPH1 recruitment and dissect whether DDX5 acts directly on the RBM3 rG4 itself or indirectly by remodelling nearby regulatory elements or co-factors. These studies will clarify the hierarchy and cooperativity among rG4 remodelling, helicase activity, and hnRNPH1-mediated exon definition.

Thus, RBM3 splicing appears to be governed by an integrated mechanism in which rG4 unfolding establishes the primary permissive state for hnRNPH1 activity, while DDX5 and kinase signalling fine-tune the efficiency and fidelity of this process (Fig. S4). This combinatorial interplay between RNA structure, helicase activity, and phosphorylation may represent a broader principle of temperature-responsive RNA regulation, helping ensure robust AS control under environmental perturbation.

Finally, the translational implications of our findings are substantial. RBM3 is a potent neuroprotective factor in models of Alzheimer’s, prion disease, and traumatic brain injury. Our results reveal that pharmacological destabilization of rG4s enhances RBM3 expression through PE skipping, offering a potential route to mimic hypothermia-induced neuroprotection without the risks associated with therapeutic cooling. More broadly, the identification of rG4s, hnRNPH1, and DDX5 as components of a temperature-sensitive splicing axis suggests new avenues for modulating RNA structure in neurodegenerative disease.

### Significance

RNA G-quadruplexes (rG4s) are well-characterized structural elements in vitro, yet their functional impact on alternative splicing in living cells has remained unclear. Here, we show that temperature-dependent remodelling of rG4 structures within RBM3 pre-mRNA governs hnRNPH1-mediated poison exon skipping, establishing RNA structural dynamics as a primary determinant of RBM3 regulation. By integrating proteomics, phosphoproteomics, and genetic screening, we further identify the RNA helicase DDX5 as a cooling-responsive auxiliary factor which is upregulated and differentially phosphorylated at 32°C and previously shown to unwind rG4s and cooperate with hnRNPH1 in splicing. These findings reveal how environmental cues such as mild hypothermia interface with RNA structure, helicase activity, and kinase signalling to modulate alternative splicing. More broadly, our work demonstrates a generalizable principle whereby temperature-driven RNA structural transitions help tune gene expression programs linked to cellular stress responses.

## Acknowledgements

The authors thank Jernej Ule and Flora lee (King’s College London) for insightful discussions. D.K. was supported by the King’s Prize Fellowship and Professor Anthony Mellows Fellowship. This research was also made possible through the support of the UK Dementia Research Institute [(UK DRI-6005 and UK DRI-6204) to M.D.R.; UK DRI Proteomics platform award (DRI-PRO2024-2) to D.K.] through UK DRI Ltd, principally funded by the Medical Research Council; the Guangdong Basic and Applied Basic Research Foundation (2023A1515111121) and the National Natural Science Foundation of China (32400453) to S.S. and J.Q.L., and Guangzhou Municipal Science and Technology Project (SL2024A04J01657) to J.Q.L.

## Author Contributions

Conceptualization: J.Q.L. and D.K.; Methodology: J.Q.L., M.D.R. and D.K.; Investigation: S.S., L.M., M.D.R., J.Q.L., and D.K.; Formal analysis: S.S., J.Q.L., and D.K.; Funding acquisition: M.D.R., J.Q.L., and D.K.; Supervision and Project administration: J.Q.L. and D.K.; Visualization: S.S., J.Q.L., and D.K.; Writing-original draft: D.K.; Writing-review and editing: S.S., L.M., M.D.R., J.Q.L., and D.K.

## Supplementary data

**Figure S1:**
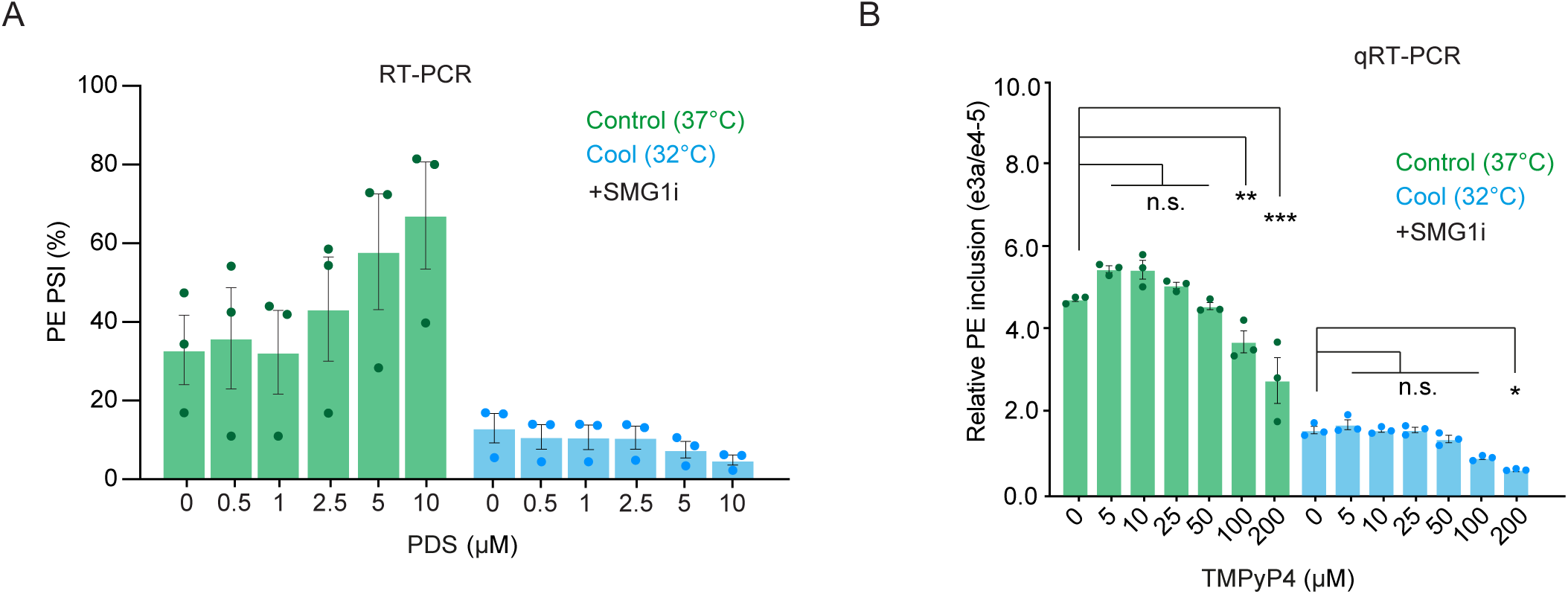
Stability of rG4 structures influence RBM3 PE skipping. (A) Graph showing the PSI values of RBM3 PE on PDS treatment which are calculated based on the intensity of PE-included (red arrows) and PE-skipped (green arrow) isoforms from RT-PCR data in Fig. 1E. (B) qRT-PCR quantifying the PSI values of RBM3 PE relative to RBM3 mRNA in HeLa at 37°C or 32°C (72h) in the presence of increasing concentration of TMPyP4 (5-200 µM). Data information: N = 3 biological replicates. Mean ± SEM; ns (not significant), *(P<0.05), **(P<0.01); ***(P<0.001); one-way ANOVA with multiple comparisons.

**Figure S2:**
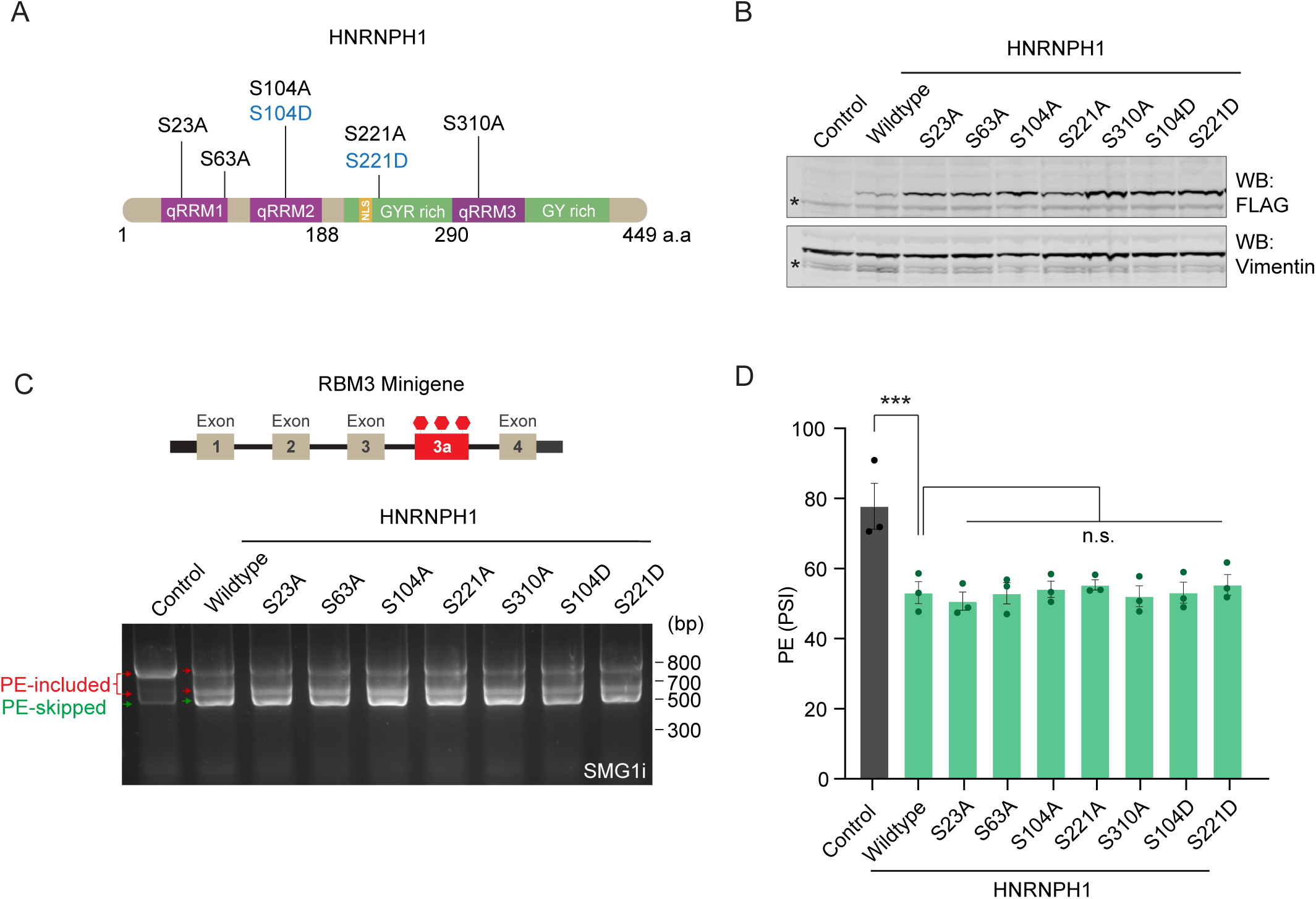
Effect of hnRNPH1 phosphorylation on RBM3 PE skipping. (A) Schematic of hnRNPH1 phosphorylation sites that were targeted for phospho-dead and phospho-active mutations. (B) Western blot from HeLa cells showing the levels of the different transfected Phosphomutants depicted in (A). GAPDH is used as loading control (C) Schematics and RT-PCR of RBM3 minigene (Exon 1–4), flanked by unique sequences (thick black bars) to distinguish it from endogenous transcripts during PCR amplification, expressed in HeLa cells which have been co-transfected with HNRNPH1 phosphomutants, and treated with SMG1 inhibitor. The PE-included (red arrows) and PE-skipped (green arrow) isoforms are shown. (D) Graph showing the PSI values of RBM3 PE which are calculated based on the intensity of PE-included (red arrows) and PE-skipped (green arrow) isoforms from (C) Data information: N = 3 biological replicates. Mean ± SEM; ns (not significant), *(P<0.05), **(P<0.01); ***(P<0.001); one-way ANOVA with multiple comparisons.

**Figure S3:**
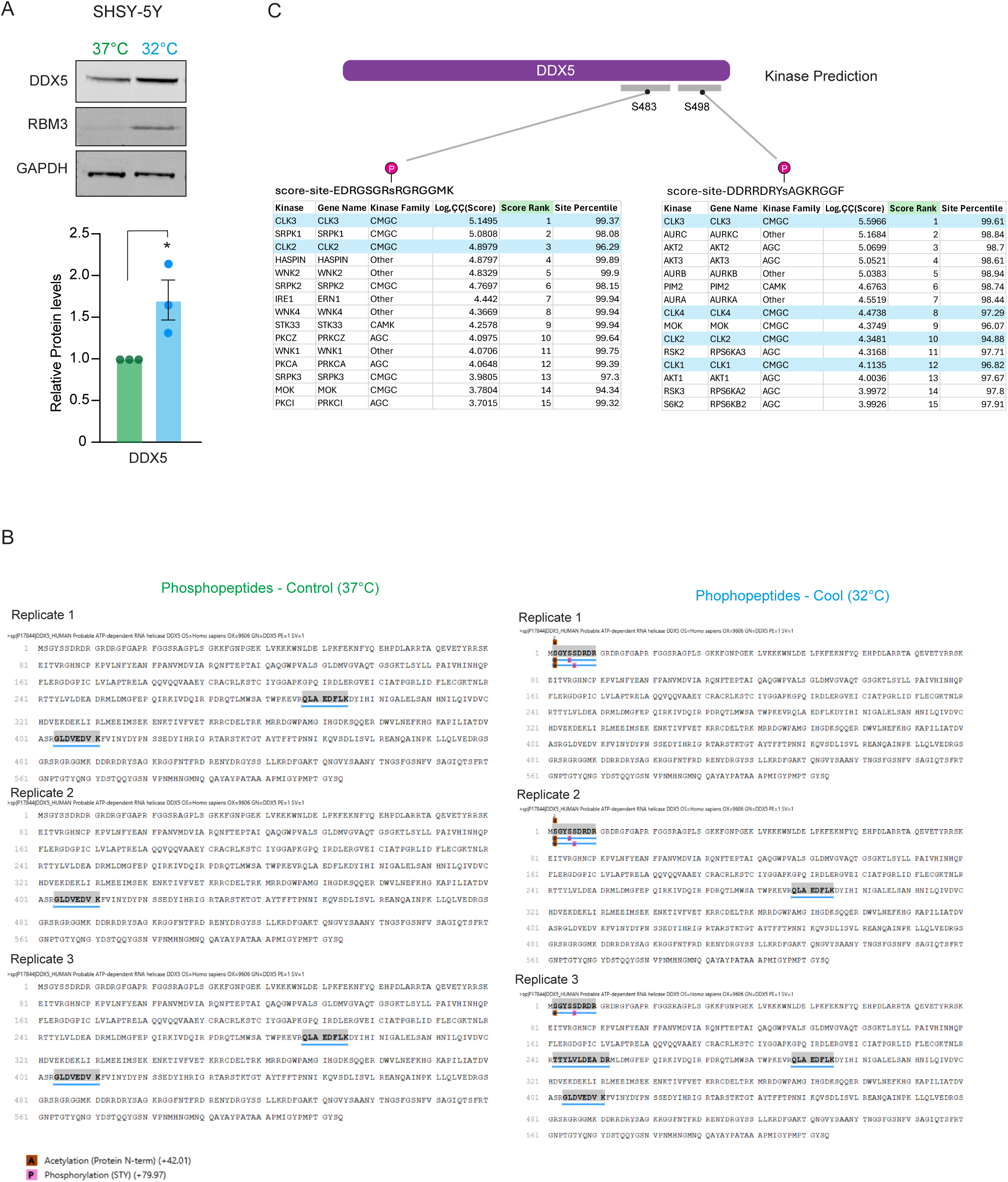
DDX5 levels and phosphorylation increase on cooling. (A) Western blot analysis of DDX5 protein levels in SHSY-5Y (neuroblastoma) cells cultured at 37°C or 32°C (72h). Representative blots are shown (left) with quantification normalized to GAPDH (right). (B) Summary of identified DDX5 phosphosites across cooling and control conditions from iPSCs. Phosphorylated residues and their corresponding peptides are mapped to either 37°C-specific, 32°C-specific, or shared phosphopeptide pools. (C) Kinase-substrate prediction analyses using *The Kinase Library* for cooling-associated DDX5 phosphopeptides. Here, for two phosphosites the top 15 predicted kinases are shown, with CLK-family kinases (CLK1/2/3/4) highlighted in blue. Multiple DDX5 phosphopeptides rank CLK enzymes among the highest-confidence motif matches, suggesting that DDX5 contains consensus sequences compatible with CLK phosphorylation. Data information: N = 3 biological replicates. Mean ± SEM; ns (not significant), *(P<0.05), **(P<0.01); ***(P<0.001); Unpaired t-test in (A)

**Figure S4:**
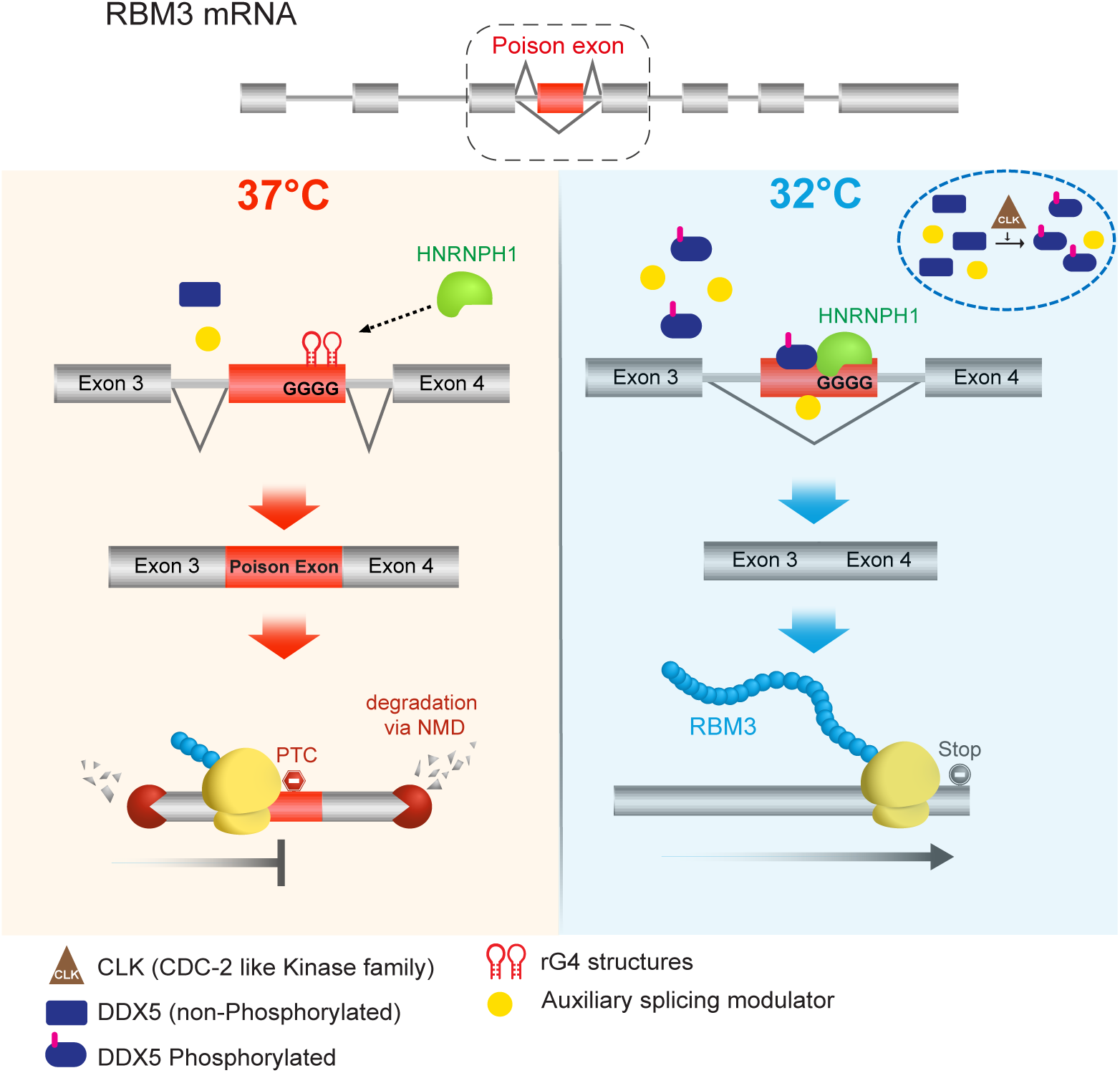
Model of rG4-dependent alternative splicing of RBM3 Poison exon. At 37°C, rG4 structures within the RBM3 PE remain folded, limiting access to hnRNPH1 and restricting PE skipping. Upon cooling to 32°C, rG4s partially unfold, enabling increased hnRNPH1 binding and promoting PE exclusion. Concurrently, DDX5 protein levels and phosphopeptide abundance increase, positioning DDX5 to cooperatively remodel rG4 or adjacent RNA structures and support hnRNPH1-mediated splicing. Predicted phosphorylation of DDX5 by CLK-family kinases provides an additional regulatory layer linking temperature-sensitive kinase activity to helicase remodelling. Together, RNA structural changes, hnRNPH1 recruitment, and DDX5 activation converge to drive enhanced RBM3 PE skipping at lower temperatures.

## Appendix Supplementary Tables

**Appendix Table S1.**
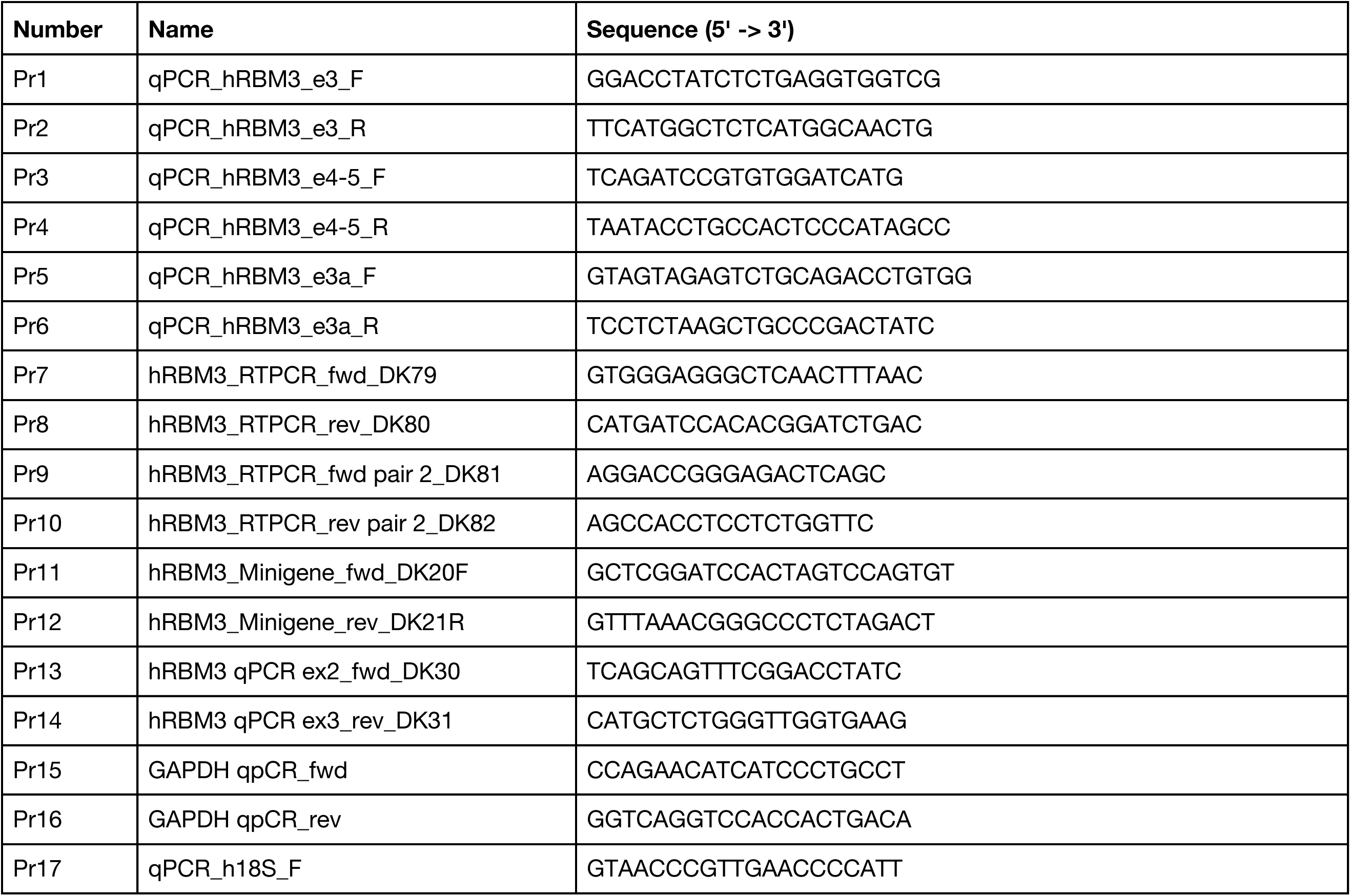
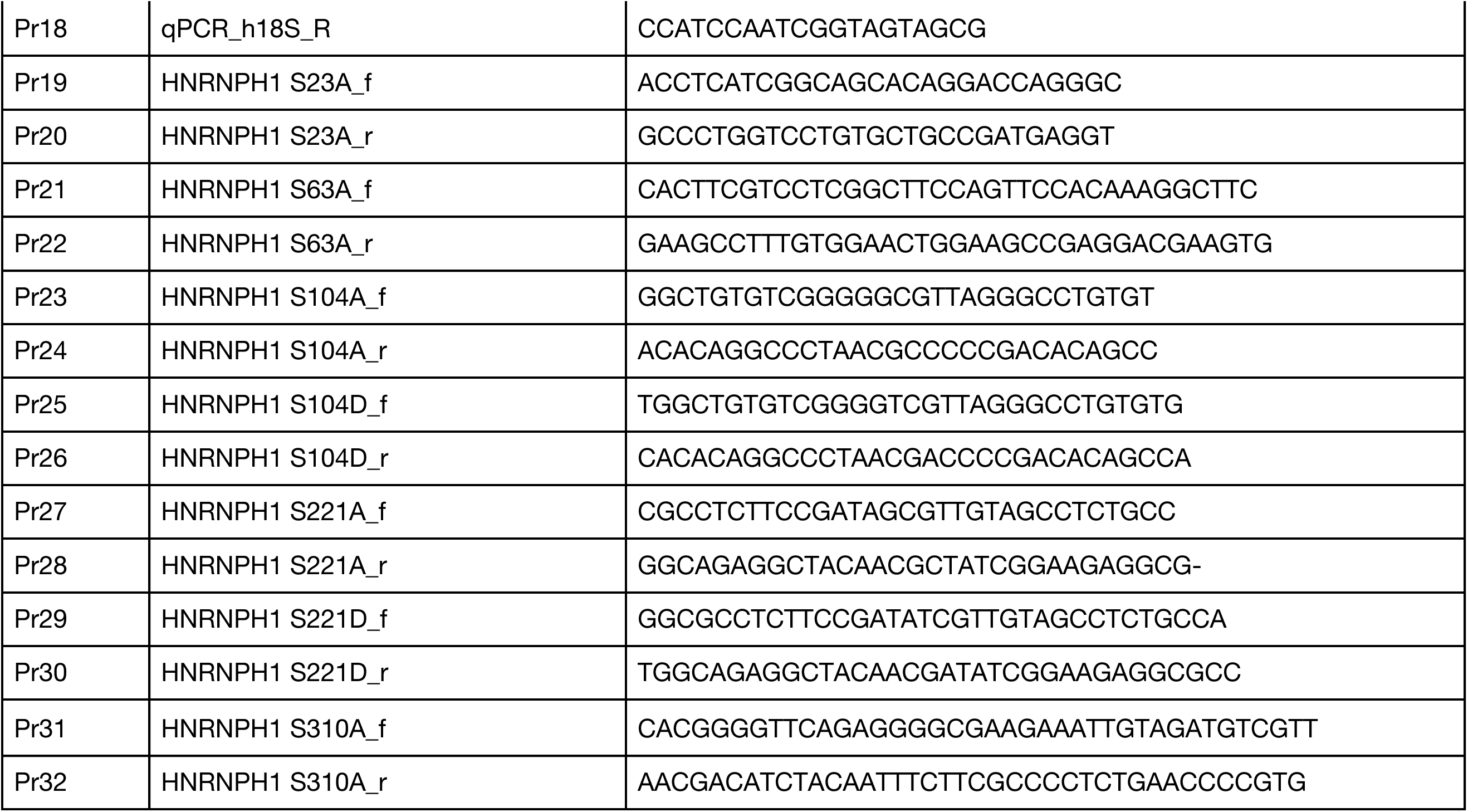
Primer (Pr) sequences used in this study (Page 2)

**Appendix Table S2.**
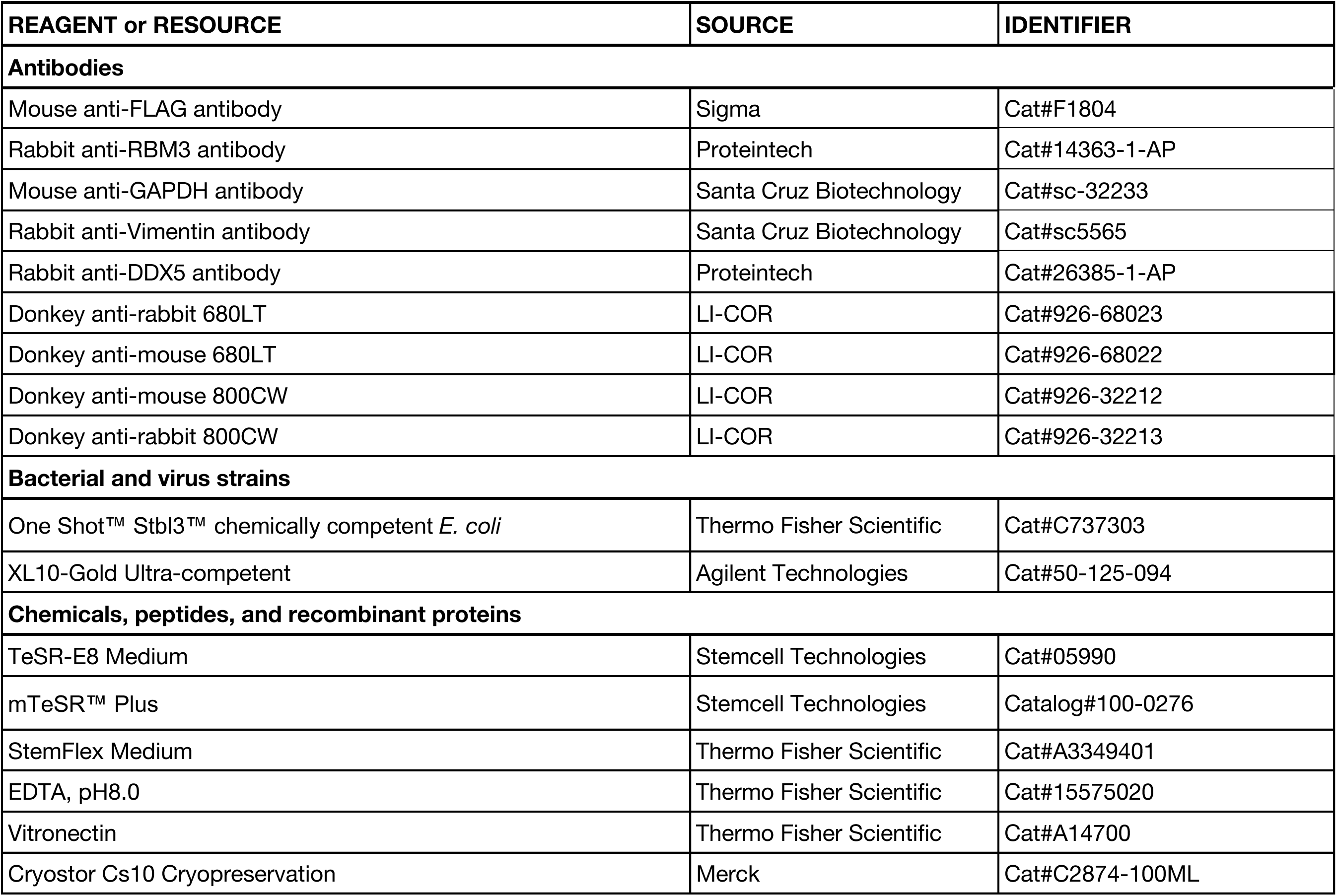
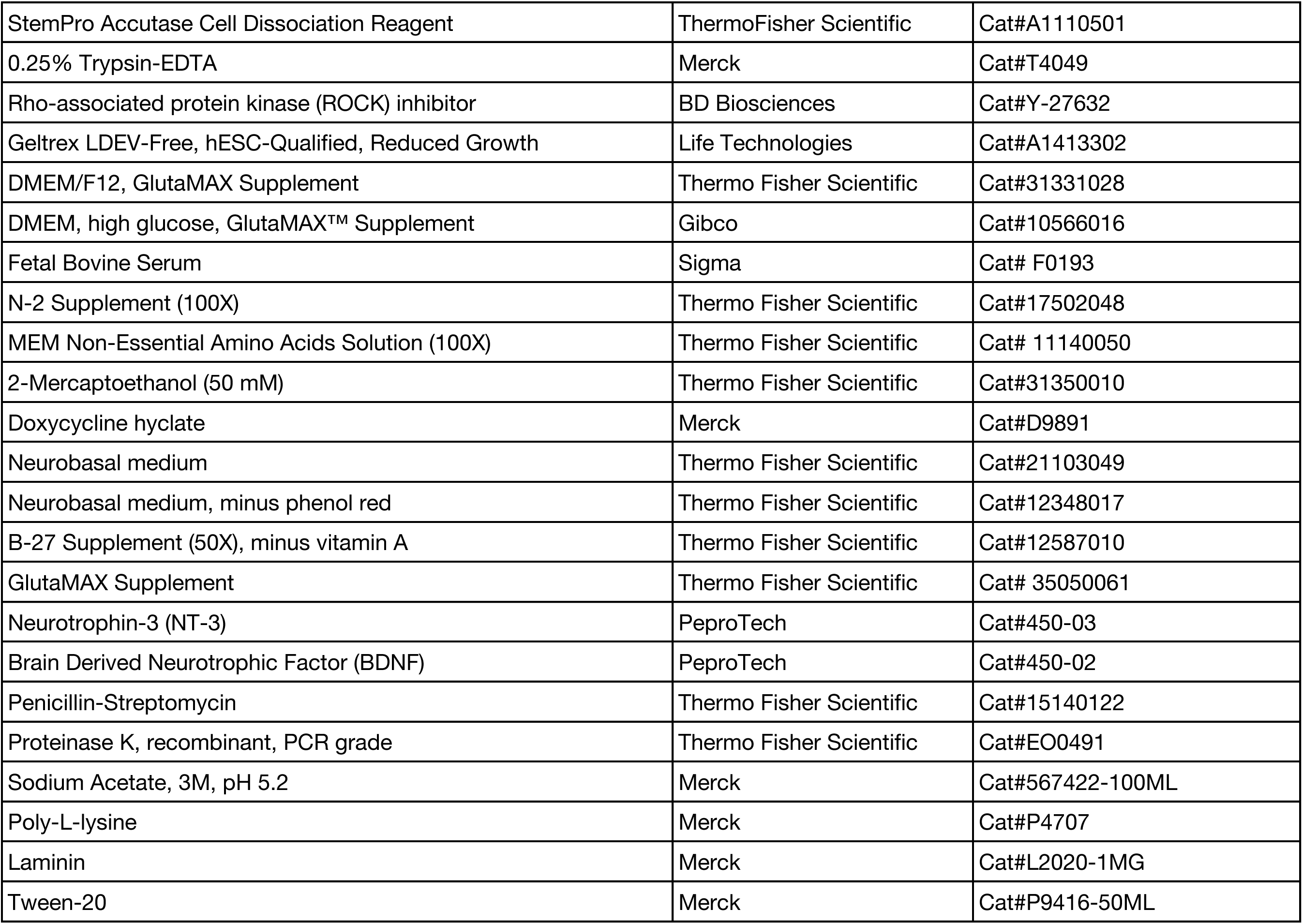
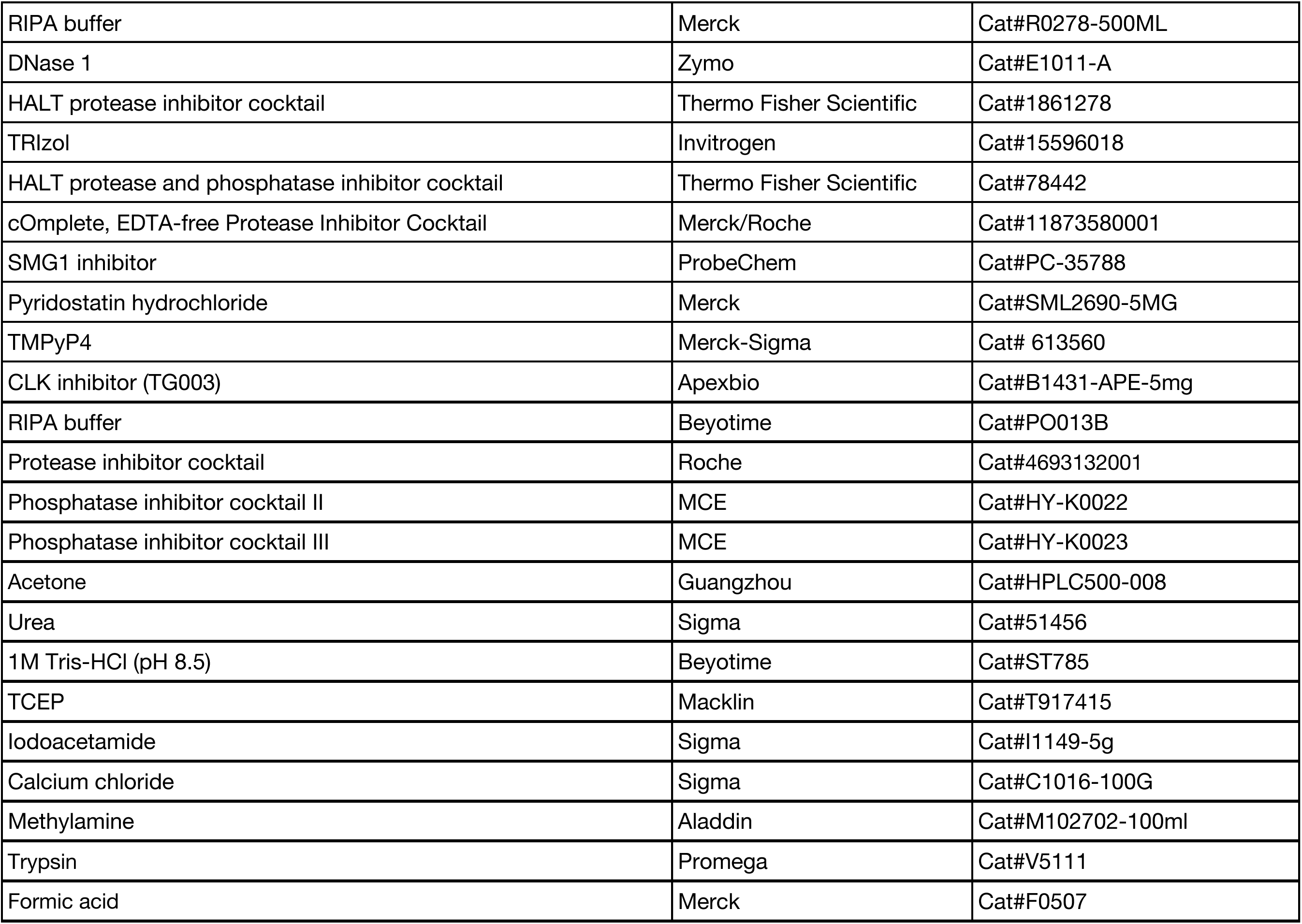
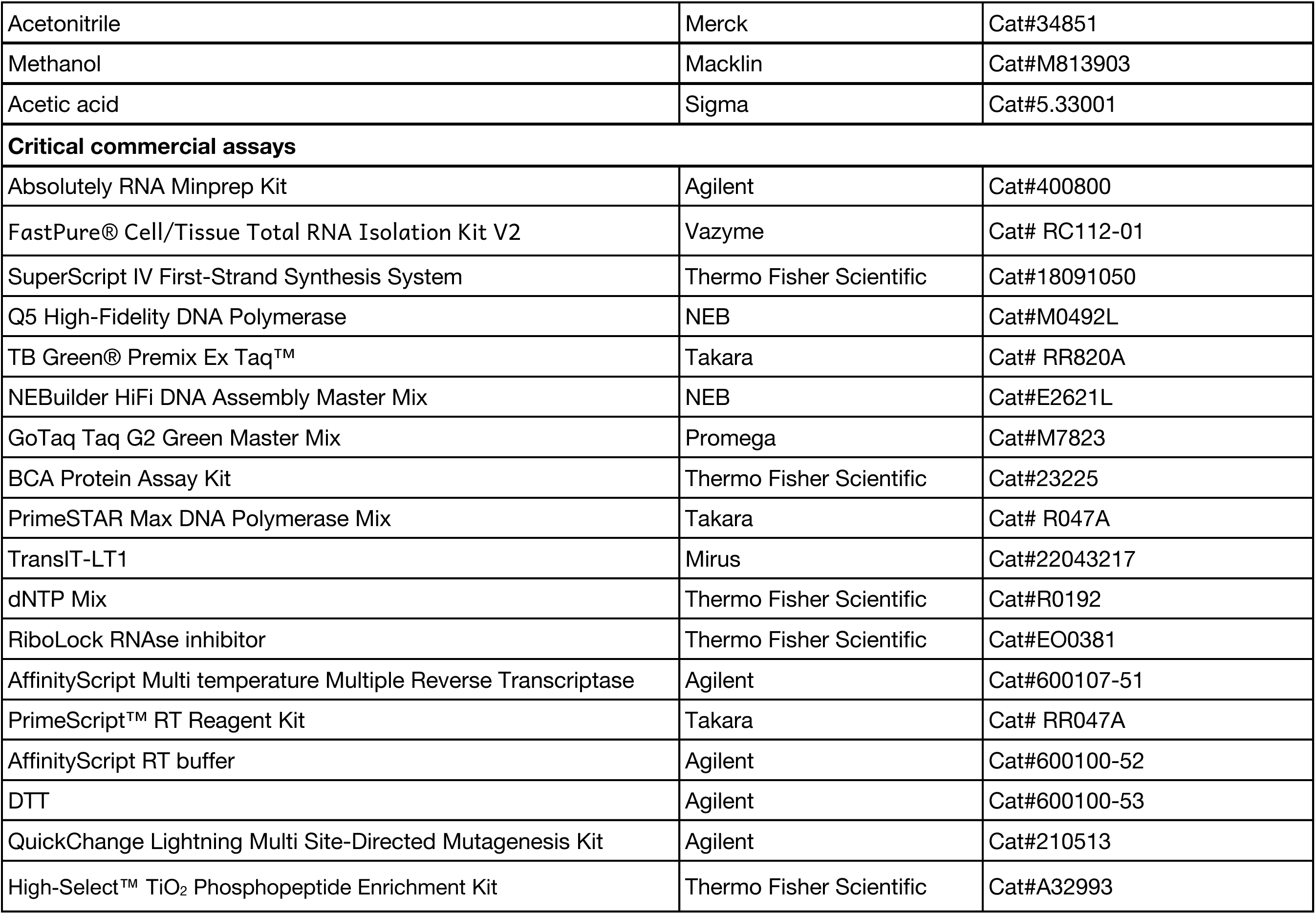
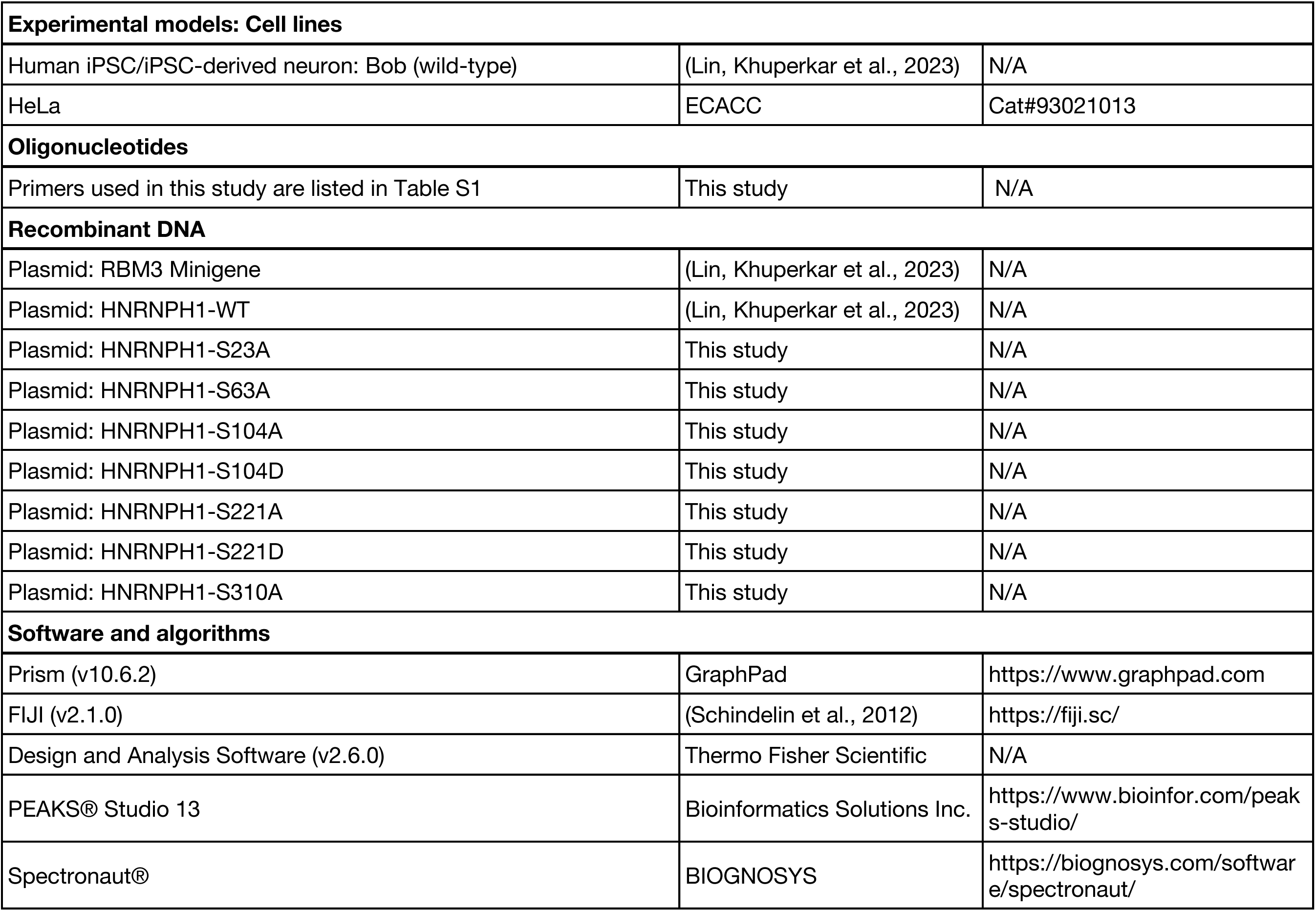
Key resources used in this study (Page 4)

**Appendix Table S3.** Top three RNA Helicases from list of 220 regulators of RBM3 identified from CRISPR screen from Lin, Khuperkar et al 2023

| id | Gene symbol | Full gene name | neg \| p-value | neg \| rank | neg \| lfc | pos \| p-value | pos \| rank |
| --- | --- | --- | --- | --- | --- | --- | --- |
| ENSG00000107201 | DDX58 | DEXD/H-box helicase 58(DDX58) | **0.0070304** | 102 | -0.28976 | **0.042293** | 2476 |
| ENSG00000089737 | DDX24 | DEAD-box helicase 24(DDX24) | **0.019821** | 215 | -0.28775 | **0.052726** | 2968 |
| ENSG00000108654 | DDX5 | microRNA 3064(MIR3064) | **0.020337** | 219 | -0.39733 | **0.73672** | 15690 |

## Material Methods

### Plasmids and oligonucleotides

All primers used in this study are listed in Appendix Table S1. Plasmids generated alongside key resources are listed in Appendix Table S2.

HNRNPH1 phosphomutants: FLAG-HNRNPH1 phosphomutants were generated using site directed mutagenesis (QuickChange Lightning Multi Site-Directed Mutagenesis Kit, Agilent) from FLAG-HNRNPH1 plasmid (described previously in Lin, Khuperkar et al., 2023) using specific primers (See Table S1). Sequences within the promoter and insert regions were confirmed by Sanger sequencing.

### HeLa cell culture and transfection

HeLa cells were grown in Dulbecco’s Modified Eagle Medium (DMEM, high glucose, GlutaMAX™ Supplement; Gibco) supplemented with 10% fetal bovine serum (Sigma) or Dulbecco’s modified Eagle’s medium (DMEM)+ Ham’s F12 (1:1) supplemented with 10% fetal calf serum (FCS), penicillin (100 IU/mL), and streptomycin (100 μg/mL) at 37°C under 5% CO₂. Plasmid DNA transfection was performed with TransIT-LT1 (Mirus) according to the manufacturer’s instructions. For HNRNPH1 overexpression experiments, 250,000 HeLa cells were seeded in a 6-well plate, and the next day cells were co-transfected with HNRNPH1 constructs and RBM3 Minigene (described previously in Lin, Khuperkar et al., 2023). After 24h, medium was changed again with fresh medium with or without 1 μM SMG1 inhibitor (Probechem, PC-35788). Cells were incubated with SMG1 inhibitor for 24h before harvesting the cells.

### Human iPSC culture and differentiation into iPSC-derived neurons (i3-neurons)

Human iPSCs with Neurogenin-2 (NGN2) transgene stably integrated into a ‘safe-harbour’ locus under doxycycline (Dox)-inducible promoter (described previously in Lin, Khuperkar et al., 2023) were maintained under feeder-free conditions in TeSR-E8 medium in a 37°C, 5% CO_2_ tissue culture incubator. They were cultured on vitronectin (3.3 µg/mL)-coated culture plates or glass-bottom dishes and fed every day with TeSR-E8 medium or every 2 days with StemFlex Medium. 0.5 mM EDTA was used for routine dissociation to maintain colony growth.

iPSCs were enzymatically detached and dissociated into single cells using Accutase and plated into GelTrex (1:100 dilution)-coated culture plates in TeSR-E8 medium supplemented with 10 µM Rho-associated protein kinase (ROCK) inhibitor. After 24 hours (day 1), TeSR-E8 medium was changed to DMEM/F12 medium with GlutaMAX, supplemented with 1x N-2 supplement, 1x Non-Essential Amino Acids, 50 nM 2-Mercaptoethanol, 100 U/mL Penicillin-Streptomycin, and 1 µg/mL Doxycycline Hyclate (Dox) (iN-1 medium). After 24 hours (day 2), the medium was replaced with the same medium as the previous day. From day 3 to day 6, the culture was fed daily with Neurobasal medium supplemented with 1x B-27 supplement (minus vitamin A), 1x GlutaMAX, 50 nM 2-Mercaptoethanol, 100 U/mL Penicillin-Streptomycin, and 1 µg/mL Dox, 10 ng/mL Neurotrophin-3 (NT-3), and 10 ng/mL Brain-derived neurotrophic factor (BDNF) (iN-2 medium). After day 6, the medium was changed every other day. The same feeding schedule as previously described was followed from day 5. I-neurons in the 32°C conditions were placed in a 5% CO_2_ incubator set to 32°C for 72h between day 15-18 post differentiation. For PE analyses, neurons were incubated with SMG1 inhibitor for 24h before collection.

### Drug treatments

#### PDS

HeLa cells were seeded and grown at 37°C. 24h post seeding, one set of cells was incubated at 32°C, while the other remained at 37°C. After 24h, the medium was replaced with fresh medium containing PDS (Merck-Sigma) at different concentrations (0, 0.5, 1, 2.5, 5 or 10 μM). Following another 24h incubation, the SMG1 inhibitor was added to a final concentration of 1 μM, and cells were harvested after an additional 24h. For i3-neurons, on day 15 post differentiation, one set of the i3-neurons was incubated at 32°C, while the other remained at 37°C. This was followed by PDS treatment with the same concentration range as described above on Day 16 (after 24h), SMG1 inhibitor incubation (1 μM) on Day 17 and harvesting on Day 18 post differentiation.

#### TMPYP4

HeLa cells were seeded and grown at 37°C. 24h post seeding, one set of cells was incubated at 32°C, while the other remained at 37°C. After 24h, the medium was replaced with fresh medium containing TMPyP4 (Merck-Sigma) at different concentrations (0, 5, 10, 25, 50, 100, or 200 nM). Following another 24h incubation, the SMG1 inhibitor was added to a final concentration of 1 μM, and cells were harvested after an additional 24h. For i3-neurons, on day 15 post differentiation, one set of the i3-neurons was changed to 32°C, while the other remained at 37°C. This was followed by TMPyP4 treatment with the same concentration range as described above on Day 16 (after 24h), SMG1 inhibitor incubation (1 μM) on Day 17 and harvesting on Day 18 post differentiation.

### CLK inhibitor (TG003)

HeLa cells were seeded and grown at 37°C. 24 hours later, one set of cells was incubated at 32°C, while the other remained at 37°C. After 24h, the medium was replaced with fresh medium containing CLK inhibitor (TG003) (Apexbio) at 10μM. Following another 24h incubation, the SMG1 inhibitor was added to a final concentration of 1 μM, and cells were harvested after an additional 24h for RNA extraction.

### RT-PCR and qRT-PCR

Total RNA of i3-neurons and HeLa cells was extracted using the Absolute RNA miniprep kit or the FastPure®Cell/Tissue Total RNA Isolation Kit V2 (Vazyme). Reverse transcription to generate cDNA was performed using the PrimeScript™ RT Reagent Kit (Takara) or AffinityScript Multiple Temperature Reverse Transcriptase. When using the latter method, 1-2 μg of isolated RNA was incubated with random hexamers (900ng) in a total volume of 67 μl in DEPC water at 65°C for 5 minutes and then for 10 minutes at room temperature. Then, the mixtures were separated into 2 parts – RT and no-RT control samples (33.5 μl each). To this 13.5 μl master mix was added to supplement 1x reverse transcriptase buffer (Agilent), 10 mM DTT (Agilent), 400 μM dNTPs (Thermo Scientific), 40 units RiboLock RNase inhibitor (Thermo Scientific) and 1 reaction equivalent AffinityScript Multiple Temperature Reverse Transcriptase (Agilent). The RT enzyme was absent in the no-RT condition. The samples were then incubated at 65°C for 1 hour followed by heat-inactivation at 75°C for 20 minutes.

For RT-PCR, 20-40 ng cDNA was amplified using GoTaq Polymerase mix or the PrimeSTAR Max DNA Polymerase Mix (Takara) using the indicated primers (Appendix Table S1). Quantification of the bands on agarose gels was done using FIJI software. Percent spliced in (PSI) indexes of RBM3 poison exon (PE) are calculated based on the intensity of PE-included (red arrows) and PE-skipped (green arrows) isoforms visualised in agarose gels:

a. One PE-included isoform is detected: (I_Inc_/L_Inc_) / (I_Inc_/L_Inc_ + I_skip_/L_skip_), or
b. Two PE-included isoforms are detected: (I_Inc1_/L_Inc1_ + I_Inc2_/L_Inc2_) / (I_Inc1_/L_Inc1_ + I_Inc2_/L_Inc2_ + I_skip_/L_skip_)

Where I_Inc_ = Intensity of PE-included isoform, L_Inc_ = length of PE-included isoform in base pairs, I_Inc_ = Intensity of PE-skipped isoform, L_skip_ = length of PE-skipped isoform in base pairs.

For qRT-PCR, cDNA from each sample (diluted 10–1,000-fold) was mixed with TB Green® Premix Ex Taq^™^ (Takara) and PCR primers in four technical replicates on 384-well plates. Reactions were run on a LightCycler® 480 Instrument II (Roche), and data were analyzed using LightCycler 480 software (Roche). Relative PE inclusion was calculated as the ratio of the RBM3 Exon 3a amplicon to either the RBM3 Exon 3 or Exon 4–5 amplicon (or their geometric mean). 18S rRNA served as the loading control for quantifying RBM3 and HNRNPH1 mRNA levels.

### Cell lysis and Western blotting

HeLa cells were harvested by scraping and washed with 1x Phosphate buffered saline (PBS). Lysis was done in 150 μl RIPA lysis buffer (Merck) per well of 6-wells plate, supplemented with 1x Halt Protease Inhibitor cocktail (Thermo Scientific), followed by sonication at 40% amplitude on ice. The extracts were then cleared by centrifugation at 15,000 x g and 4°C for 15 minutes. Protein concentrations of lysates were determined by BCA assay following the manufacturer’s instruction. Samples were diluted with Laemmli protein sample buffer with 100 mM DTT. 20 μg protein was loaded into each well on a 4–12% Bis-Tris polyacrylamide gels (NuPAGE) followed by western blotting and ran at 150 V. Gels were dry blotted onto nitrocellulose membranes on an iBlot2 gel transfer system (Invitrogen). Membranes were blocked with 5% non-fat milk in 1x TBS-T for 1 h rotating at room temperature. The primary antibody solution was incubated overnight at 4°C while rotating. The next day, membranes were washed three times with 1x TBS-T, then incubated for 1 h in secondary antibody solution (1:10,000 in 5% non-fat milk in 1x TBS-T) and washed three times with 1x TBS-T before imaging on LI-COR Odyssey CLx. Primary and secondary antibodies were used in the following concentrations: Rabbit anti-RBM3 (1:1500), mouse anti-GAPDH (1:3000), Rabbit anti-DDX5 (1:3000), Rabbit anti-Vimentin (1:2000), mouse anti-FLAG (1:3000), donkey anti-mouse 680LT (1:10000), donkey anti-rabbit 800CW (1:10000) and donkey anti-mouse 800CW (1:10000).

### hnRNPH1 phosphosites prediction

Putative phosphorylation sites within full-length human HNRNPH1 (UniProt ID: Q00464) were identified using the PhosphoSitePlus database (Hornbeck et al., 2015). Candidate residues were shortlisted according to the following criteria: (i) distribution across all three RNA recognition motifs (RRMs) to ensure domain-wide coverage, (ii) evolutionary conservation between human and mouse sequences, and (iii) high-confidence phosphorylation scores reported in the database. Sites fulfilling these criteria were selected for subsequent site-directed mutagenesis to generate phospho-deficient (serine to alanine) and phospho-mimetic (serine to aspartate) hnRNPH1 constructs for functional assays.

### Total proteomics and Phospho-proteomics

The lysis buffer, composed of RIPA (Beyotime) supplemented with 1X protease inhibitor cocktail (Roche), 1X phosphatase inhibitor cocktail II (MCE), and 1X phosphatase inhibitor cocktail III (MCE), was directly added to the culture dish of iPSC cells. Prior to lysis, the cells were washed once with 1× Dulbecco’s phosphate-buffered saline (DPBS). The cells were then harvested using a cell scraper. The resulting cell lysate was transferred to a centrifuge tube. Four volumes of pre-chilled acetone (-20°C) were added to the lysate, followed by vortexing and incubation at - 20°C overnight. After incubation, the sample was centrifuged at 12,000 × g and 4°C for 10 minutes. The supernatant was discarded, and the pellet was washed once with a fresh volume of pre-chilled acetone. The pellet was then air-dried in a biosafety cabinet with the tube lids open at room temperature to evaporate residual acetone. The proteins were solubilized in a freshly prepared solution of 8M urea (Sigma) and 100 mM Tris (pH 8.5) (Beyotime). Protein concentration after precipitation was determined by a BCA assay (Thermo) to guide subsequent trypsin usage. For reduction, TCEP (Macklin) was added to a final concentration of 5 mM, and the sample was incubated at room temperature for 20 minutes. This was followed by alkylation through the addition of iodoacetamide (Sigma) to a final concentration of 10 mM, with incubation in the dark at room temperature for 15 minutes. The sample was then diluted four-fold with 100 mM Tris (pH 8.5) to reduce the urea concentration.

CaCl₂ was added to a final concentration of 1 mM, and methylamine was added to 20 mM to minimize carbamylation. Trypsin (Promega) was added at a 1:100 (w/w) enzyme-to-protein ratio. Digestion was carried out in the dark at 37°C overnight (∼16 h). The tryptic digestion was stopped by acidifying the sample with formic acid (Merck) to a final concentration of 0.5%. For desalting, a large C18 column (Waters) was used. The column was conditioned with 1 mL of methanol, washed with 1 mL of 80% acetonitrile (Merck) (in 0.5% acetic acid (Sigma)), and equilibrated with 1 mL of 0.5% acetic acid. The acidified sample was loaded onto the column, which was then washed three times with 1 mL of 0.5% acetic acid. Peptides were eluted by applying 500 µL of 80% acetonitrile, and the same eluent was passed through the column a second time into the same tube. The peptide eluate was dried in a SpeedVac vacuum concentrator at 35°C for approximately 2h. The resulting peptide powder was stored at -80°C. The peptide sample was subsequently split: one portion was used for total proteome analysis, and the other portion was subjected to phosphopeptide enrichment using the High-Select™ TiO2 Phosphopeptide Enrichment Kit (Thermo Fisher Scientific) according to the manufacturer’s instructions. Prior to LC-MS/MS analysis, the peptides were reconstituted in 0.1% formic acid. All LC-MS/MS analyses were performed on a timsTOF HT mass spectrometer coupled with a nanoElute 2 HPLC system (Bruker) at HKUST(GZ) Biosciences Central Research Facility (BioCRF). For total proteome analysis, the samples were analyzed in data-independent acquisition (DIA) mode, and the resulting data were processed using Spectronaut software for label-free quantification. For phosphoproteome analysis, the enriched phosphopeptides were analyzed in data-dependent acquisition (DDA) mode. The DDA data from the phosphoproteome were processed and analyzed using PEAKS Studio software for phosphopeptide and phosphosite identification.

### Kinase prediction analyses

Kinase prediction analysis for DDX5 was performed using the PhosphoSitePlus Kinase Library tool (https://kinase-library.phosphosite.org/kinase-library/) (Johnson et al., 2023). All experimentally reported human DDX5 phosphopeptides listed in PhosphoSitePlus were extracted and individually submitted to the “Scan Substrate” module using default settings. For each phosphopeptide, the algorithm scores motif similarity against curated kinase recognition motifs and returns a ranked list of candidate kinases. Rankings for the top predicted kinases were recorded for each DDX5 phosphosite. Across multiple independent DDX5 phosphopeptides, CLK-family kinases (CLK1, CLK2, CLK3 and CLK4) consistently appeared among the highest-scoring motif matches, indicating that DDX5 contains sequence motifs compatible with CLK phosphorylation (Fig. S3).

## Notes

### Competing Interest Statement

The authors have declared no competing interest.

### Summary of Updates

Author contributions have been updated with more information on Funding acquisition and project administration contributions

